# GWAS SVatalog: a visualization tool to aid fine-mapping of GWAS loci with structural variations

**DOI:** 10.1101/2025.09.03.674075

**Authors:** Shalvi Chirmade, Zhuozhi Wang, Scott Mastromatteo, Eric Sanders, Bhooma Thiruvahindrapuram, Thomas Nalpathamkalam, Giovanna Pellecchia, Fan Lin, Katherine Keenan, Rohan V Patel, Wilson WL Sung, Delnaz Roshandel, Joe Whitney, Sana Allana, Julie Avolio, Paul Eckford, Felix Ratjen, Lisa J Strug

## Abstract

**Background:** Genome-wide association studies (GWAS) have been successful in identifying single nucleotide polymorphisms (SNPs) associated with phenotypic traits. However, SNPs form an incomplete set of variation across the genome and since a large percentage of GWAS-significant SNPs lie in non-coding regions, their impact on a given trait is difficult to decipher. Recognizing whether these SNPs are tagging other polymorphisms, like structural variations (SV), is an important step towards understanding the putative causal variation at GWAS loci.

**Results:** Here, we develop GWAS SVatalog (https://svatalog.research.sickkids.ca/), a novel open-source web tool that computes and visualizes linkage disequilibrium (LD) between SVs and GWAS-associated SNPs throughout the human genome. The tool combines GWAS Catalog’s SNP-trait association data across 14,479 phenotypes with LD statistics calculated between 35,732 SVs and 116,870 SNPs identified in 101 whole-genome long-read sequences. We use GWAS SVatalog to identify SVs that may explain GWAS loci for iron levels, refractive error, and Alzheimer’s disease, where previously SNPs were unable to provide a causal explanation.

**Conclusions:** GWAS SVatalog advances the fine-mapping of GWAS loci with structural variations, enabling researchers to associate 35,732 common SVs with 14,479 phenotypes, accelerating the understanding of disease etiology.

## Introduction

Genome-wide association studies (GWAS) have been successful in identifying genetic variants associated with human traits (Visscher *et al*., 2017; Uffelmann *et al*., 2021). GWAS identification of a protein-coding variant informs understanding of the mechanism by which an associated variant contributes to the analyzed trait, however, 93% of GWAS associated loci (GAL) identified lie in non-coding regions of the genome (Maurano *et al*., 2012). Such regions frequently harbor regulatory features but the mechanism by which the associated locus impacts the trait is not immediately obvious. These features may be in proximity to the affected gene or farther away where the 3D structure of the chromosome contributes to regulation (Orozco *et al*., 2022). GWAS typically use single nucleotide polymorphisms (SNPs) (Visscher *et al*., 2017; Uffelmann *et al*., 2021), which are straightforward to study for association on a large scale as they are mostly biallelic and highly quantifiable using microarray and/or short-read whole-genome sequencing (WGS). However, SNPs account for an incomplete proportion of genetic variation and phenotypic heritability (Zarrei *et al*., 2015), and it is unknown how often they tag other polymorphisms, such as structural variants, that may be the cause of the association signal.

A structural variant (SV) is a large genetic polymorphism, typically defined to range from 50 bp to several megabases and includes insertions, deletions, duplications and inversions. SVs can have a pronounced effect on gene regulation with downstream phenotypic consequences (Chiang *et al*., 2017; Vialle *et al*., 2022; Trost *et al*., 2022). Due to their size and complexity, identification and genotyping of SVs is difficult using short-read sequencing technologies. Short reads often fail to map correctly near a large SV due to reference bias (Meynert *et al*., 2014). Due to the size of short reads, most SVs are not found within a single read; their identification requires overlapping short reads to infer an SV signature (Mahmoud *et al*., 2019). Databases that catalog SVs from technologies that include short reads exist, such as the Database for Genomic Variants (DGV) (MacDonald *et al*., 2014) and the Genome Aggregation Database (gnomAD) (Chen *et al*., 2022). These are remarkable public resources used extensively by researchers, but the content of these resources is limited by the technology that is used. Since the advancement of long-read sequencing technologies, the accuracy in detecting complex SVs has increased dramatically due to longer read lengths, improved *de novo* assemblies and new SV-specific callers (Pacific Biosciences 2018). However, despite these advancements, SV detection, characterization, boundaries and allele-differentiation remains imperfect. Nevertheless, we can substantially augment these resources to aid in fine-mapping of GAL by calling common SVs from long-read WGS and pre-computing their linkage disequilibrium (LD) with GWAS SNPs, which we demonstrate here with our new tool, GWAS SVatalog.

SVs are important to genotype accurately as they have been previously shown by fine-mapping to explain GWAS signals. A motivating example from our own work is a GWAS of intestinal obstruction at birth in cystic fibrosis (CF), where we identified a suggestive locus at chr7q35 (Gong *et al*., 2019). Fine-mapping of this region using linked-read WGS uncovered a large 20-kb deletion SV in high LD with the GWAS-suggestive SNPs (Mastromatteo *et al*., 2023). We show that this SV is an eQTL for serine protease 2 (*PRSS2*) which encodes the digestive enzyme anionic trypsinogen (Mastromatteo *et al*., 2023), and is the putative cause for the GWAS locus. Discovery of SVs as the cause of GAL has been reported by others as well (Fritsche *et al*., 2008; Trost *et al*., 2022). For example, in a GWAS investigating age-related macular degeneration (AMD), a study first identified rs10490924 as associated with disease risk (Rivera *et al*., 2005) and this was replicated in multiple independent studies (Jakobsdottir *et al*., 2005; Fritsche *et al*., 2008; Micklisch *et al*., 2017). Fine-mapping of the GWAS locus identified an SV in high LD with rs10490924 located in the 3’UTR of *ARMS2* (Fritsche *et al*., 2008). This SV was shown to remove the poly-A tail of the transcript and affect expression of the protein (Micklisch *et al*., 2017), ultimately influencing the risk of AMD.

Many GWASs have identified trait-associated SNPs with no reported cause (Chen *et al*., 2021; Stachowska *et al*., 2022; Lee *et al*., 2023). Here we create a population level catalog of common SVs called from long-read WGS. The SV calls were derived from sequencing the DNA of 101 individuals enrolled in the CF Canada-SickKids Program in Individualized Therapy (CFIT) (Eckford *et al*., 2019), using both PacBio continuous long-read (CLR) and 10X Genomics (10XG) linked-read technologies. Using two separate sequencing platforms is rare for SV catalogs and has the benefit of increasing the reliability and robustness of the calls. Current publicly available long-read databases are not population-based and contain diverse ethnicities with small sample sizes (Liao *et al*., 2023; Logsdon *et al*., 2024), minimizing the reliability of LD required for this study’s analysis. We show here that the allele frequencies (AF) across the genome of individuals with CF are comparable to individuals of European origin in the general population. The curated list of SVs cataloged here has been benchmarked against well-studied genomes and annotated variant databases. Using our sequences, we compute the LD between 35,732 SVs and 116,870 GWAS-associated SNPs provided in the GWAS Catalog (Sollis *et al*., 2023), thereby evaluating association between common SVs and 14,479 human traits. We aid fine-mapping and functional follow-up for the research community by cataloguing and visualizing the LD in our novel web tool, GWAS SVatalog (https://svatalog.research.sickkids.ca/), as part of the LocusFocus (Panjwani *et al*., 2020) suite of software tools.

## Materials and Methods

### Sample cohort and DNA extraction

CFIT is a collaborative project and biospecimen repository of participants with CF to aid in advancing personalized CF therapies (Eckford *et al*., 2019). CFIT recruited and obtained whole blood from 101 Canadians with CF. Methods for recruitment, biospecimen collection and data generation are described in Eckford *et al*. 2019 (Eckford *et al*., 2019). Here we use the resulting sequencing data.

### Library preparation & sequencing

#### PacBio CLR

Library preparation and DNA extraction details are provided in Eckford *et al*. 2019 (Eckford *et al*., 2019). PacBio Sequel I and II were used to carry out the long-read sequencing on 34 and 67 samples respectively. Although the samples were sequenced on two different platforms, we did not find a significant difference between their average read lengths. The average coverage for both machines is 50x and 76x respectively.

#### 10XG Linked-Reads

Library preparation and DNA extractions details are provided in Eckford *et al*. 2019 (Eckford *et al*., 2019). Illumina HiSeq X was used to carry out the paired-end sequencing of approximately 150 bases in length at 30x coverage.

### Variant calling

For each sample, we used a combination of callers for each sequencing platform previously shown to achieve the best results (Coutelier et al. 2022; Mahmoud et al. 2024). To align and call SVs from the PacBio CLR sequences, the combination of pbmm2 1.1.0 (Pacific Biosciences 2020) + pbsv 2.2.2 (Pacific Biosciences 2018) and NGMLR 0.2.8 (Sedlazeck *et al*., 2018) + Sniffles 1.0.11 (Sedlazeck *et al*., 2018) was used. For the 10XG sequences, Long Ranger 2.2.2 (10X Genomics 2020) was used for sequence alignment, and SV calling was conducted on Long Ranger, CNVnator 0.4 (Abyzov *et al*., 2011), ERDs 1.1 (Zhu *et al*., 2012) and Manta 1.6.0 (Chen *et al*., 2016). All software was run using GRCh38 including its alternative contigs as the reference genome at default settings except for the --min_support parameter on Sniffles, which was set to five minimum reads. Samples run on Sequel I had their minimum read threshold reduced to five reads, instead of the default of ten due to lower coverage. SNPs were called from the 10XG sequences in Long Ranger using the Haplotype Caller function in GATK 4.0.0.0 (Van der Auwera and O’Connor, 2020) with default parameters.

### Comparison of allele frequencies between the CF cohort and EUR population from 1000 Genomes Phase 3

Using the list of high-confidence SNPs taken from Illumina Omni 2.5-8 v1.5 (https://webdata.illumina.com/downloads/productfiles/humanomni25/v1-5/infinium-omni2-5-8v1-5-a1-manifest-file-csv.zip), we compared the AFs of these SNPs between our CF cohort and the EUR population from 1000 Genomes Phase 3 (1000 Genomes Project Consortium *et al*., 2015) using Fisher’s Exact Test in PLINK 1.90beta3a (Purcell *et al*., 2007). Aside from the CF-causing variants on chromosome 7 and the LD block in which they lie, the remainder of the genome does not differ significantly in AF from the 1000 Genomes European population (Supplementary Figure 1).

### Merging SV calls

For the PacBio CLR sequences, pbsv and Sniffles were utilized while the 10XG sequences used Long Ranger, CNVnator, ERDs, and Manta. SV calls were finalized after three merging steps: within the PacBio CLR sequences, within the 10XG sequences, and between the two sequencing platforms (PacBio CLR and 10XG). In all steps, a 50% reciprocal overlap rule was used for deletions, inversions and duplications where the SV boundaries are finalized as the average breakpoints from the constituting calls. For insertions the criteria were slightly different, where we made sure the breakpoints were within 1,000 bp and the length of the SV within 50% of one another. The rendered call may not have the exact coordinates of the polymorphism itself but is representative of the region indicating a prominent SV.

The first step commenced by filtering pbsv calls with similar breakpoints in repeat regions. This was done to avoid duplication but does not take into account varying alleles in the individual. The pbsv calls are used as an anchor while the calls made by Sniffles were used as supporting evidence. When both pbsv and Sniffles identify the same SV, the Sniffles call would “tag” the pbsv coordinates and only the pbsv call (and genotype) would be used moving forward. This process of tagging is used to help eliminate repetition. As pbsv uses consensus sequences from supporting reads to determine SV boundaries, we chose to use its output as the primary result. However, all non-tagged calls (unique calls), made by each software are also retained moving forward. In the second step, Manta calls were used as the anchor and the supporting calls were sequentially added in the order of CNVnator, ERDs followed by Long Ranger. Only deletion calls were used from Long Ranger as the software does not identify insertions and the calls made for duplications and inversions were disproportionate in comparison to the other software. The last merging step follows the technique used in Audano *et al*. 2019 for merging SV calls across both sequencing platforms. The boundaries defining the SV are the ones from the PacBio CLR calls when present in both sequencing platforms.

To complete the merging step, SVs across all samples were consolidated to create a comprehensive SV database. SVs with overlapping coordinates were only included into the final data set if at least one call was greater than 50 bp in length. Each merging step was carried out using bcftools merge 1.20 (Danecek *et al*., 2021).

Lastly, SV calls which appear in fewer than three participants were removed from the database for the purpose of maintaining GA4GH principles (Rehm *et al*., 2021), bringing the total SVs in the database down from 129,485 to 87,183 SVs. The overview of the pipeline implemented can be seen in Figure 1.

**Figure 1:**
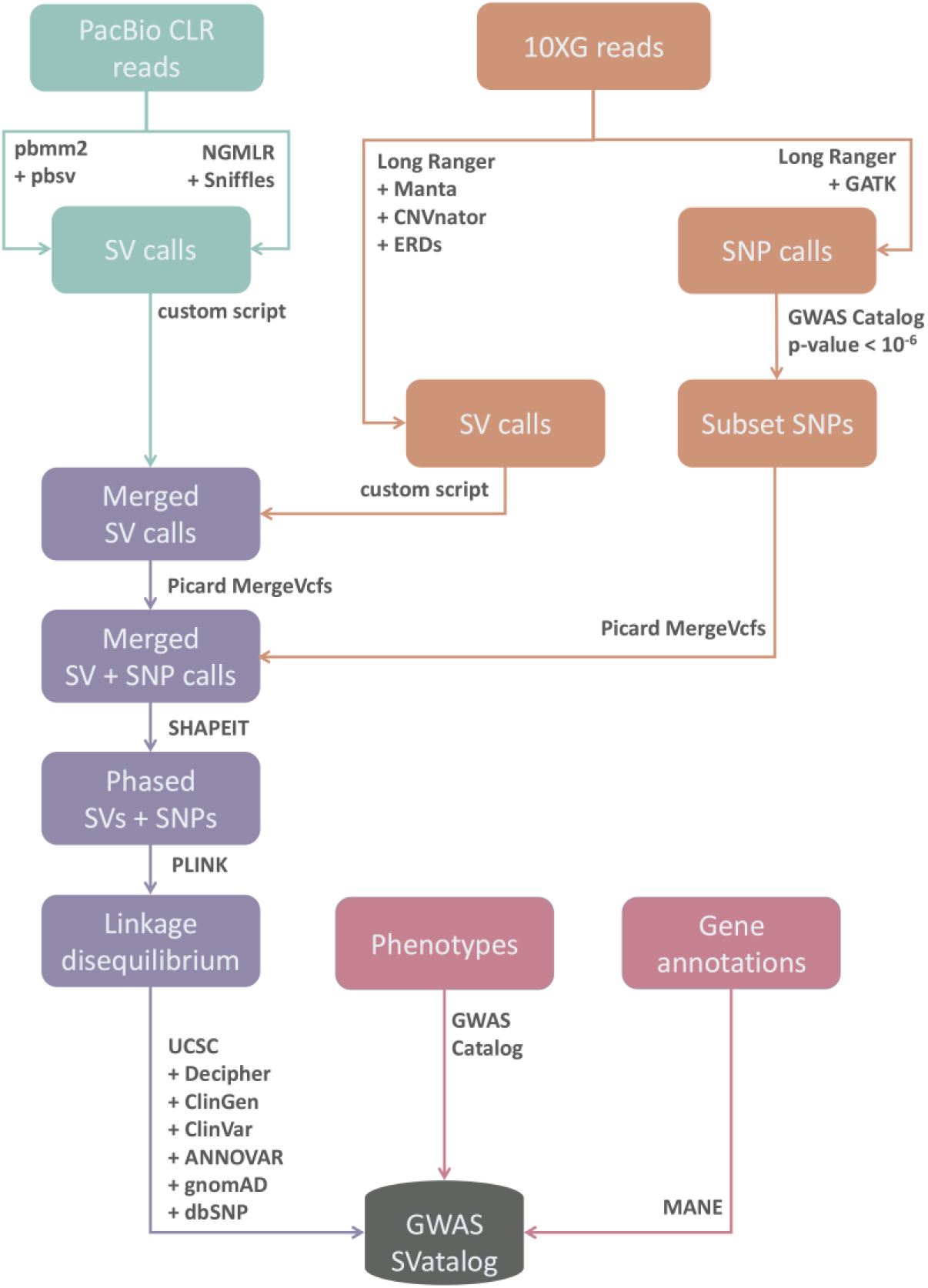
A detailed flowchart of the methods and software used in the pipeline for calling and merging SVs and SNPs across samples. We followed the implementation described by Mahmoud *et al*. 2024 for the combination of software to call SVs using PacBio long-reads, Coutelier *et al*. 2022 for the combination of software to call SVs using 10XG linked-reads, and Audano *et al*. 2019 for the merging technique of SV calls between the sequencing platforms.

### Reliability of SV calls

The SV database was validated based on SV boundaries and non-reference allele frequency (NAF). Truvari (English et al. 2022) was used to compare SV calls with three public SV datasets (Zook et al. 2016; Audano et al. 2019; Beyter et al. 2021) using the parameters `truvari bench --refdist 500 -- pctsize 0.7 --pctoval 0.5`. The SV dataset by gnomAD v4.1 (Chen et al. 2024) was used to compare SV boundaries and NAF to this study. We calculated the concordance correlation coefficient (CCC) in the two datasets and reported the confidence intervals (CI). In addition, our SV genotypes were validated by using HG002 sequences for both 10XG and PacBio CLR from Genome In A Bottle (GIAB) (https://github.com/genome-in-a-bottle/giab_data_indexes). These sequences were put through the same SV calling pipeline as our samples and the genotypes were compared with the draft release HG002 benchmark file from GIAB (https://ftp-trace.ncbi.nlm.nih.gov/ReferenceSamples/giab/data/AshkenazimTrio/analysis/NIST_HG002_DraftBenchmark_defrabbV0.019-20241113/GRCh38_HG2-T2TQ100-V1.1_stvar.vcf.gz). The comparison was carried out using the same Truvari parameters outlined above which provides genotype matching statistics.

### Linkage disequilibrium statistics

LD refers to the nondependent inheritance of alleles where, in this study, we focus on the relationship between SNPs and SVs. The two statistics, r^2^ and D’, are calculated and indexed in the tool to capture LD between the polymorphisms. In order to begin calculating the LD statistics, Picard MergeVcfs 2.26.8 (“GitHub - broadinstitute/picard: A set of command line tools (in Java) for manipulating high-throughput sequencing (HTS) data and formats such as SAM/BAM/CRAM and VCF.”) is used to consolidate the SV calls from both sequencing platforms and SNP calls from the 10XG sequences. The SNP calls in the 101 samples are filtered by selecting the SNPs reported in GWAS Catalog with a p-value < 10^-6^. Next, SHAPEIT4.2.0 (Delaneau *et al*., 2019) is used to estimate phase on the variant calls as the SNP calls from PacBio CLR sequences are presumed to be error prone. Phasing error rates using SHAPEIT4 have been shown by previous researchers to be low for both PacBio (0.23%) and 10XG (0.07%) (Delaneau *et al*., 2019). LD statistics are calculated for GWAS Catalog SNPs within 1 Mb of each SV by PLINK 1.90b3x.

### Annotation of SNPs and SVs

The custom annotation pipelines for SNPs and SVs were developed at The Centre for Applied Genomics (TCAG) at The Hospital for Sick Children and have been implemented in several studies (Chan *et al*., 2022; Trost *et al*., 2022). The SV calls were annotated using a custom CNV/SV annotation pipeline based on UCSC tracks (Nassar *et al*., 2023), gnomAD v4.1, Decipher v11.25, ClinVar/ClinGen (Landrum *et al*., 2020) portals and DGV gold standards (MacDonald *et al*., 2014). The SNPs were likewise annotated using a custom small variant annotation pipeline based on ANNOVAR v2019Oct24 (Wang *et al*., 2010), gnomAD v3.1 (Gudmundsson *et al*., 2022) and dbSNP v138 (Sherry *et al*., 2001).

### Merging with public data

Merging our LD statistics with GWAS Catalog uses the genomic position of SNPs, their reference and alternate allele. Supplementary Figure 2 shows the details of each merging step and how the data was incorporated into the database used by GWAS SVatalog. GWAS Catalog v1.0-associations_e108 was obtained from https://www.ebi.ac.uk/gwas/docs/file-downloads. MANE Select (Morales *et al*., 2022) from https://ftp.ncbi.nlm.nih.gov/refseq/MANE/MANE_human/release_1.0/MANE.GRCh38.v1.0.ensembl_genomic.gtf.gz was used to visualize genomic location in GWAS SVatalog. The curated SV database along with SNP calls and their LD statistics (r^2^ and D’) data is publicly accessible in a Zenodo repository at https://zenodo.org/records/13367574. The database indexes 35,732 SVs associated with phenotypes from the GWAS Catalog.

### GWAS SVatalog: web application development

GWAS SVatalog (https://svatalog.research.sickkids.ca/) is a web-based visualization tool built using Plotly Dash and Python (Figure 2). It is part of the LocusFocus suite of software tools (https://locusfocus.research.sickkids.ca/) (Panjwani *et al*., 2020) specializing in data integration for GWAS follow-up. GWAS SVatalog allows users to visualize the LD between 35,732 common SVs and 116,870 GAL found for 14,479 phenotypes from the GWAS Catalog. For a detailed description on how to use the tool, visit the GWAS SVatalog’s documentation (https://gwas-svatalog-docs.readthedocs.io/en/latest/index.html).

**Figure 2:**
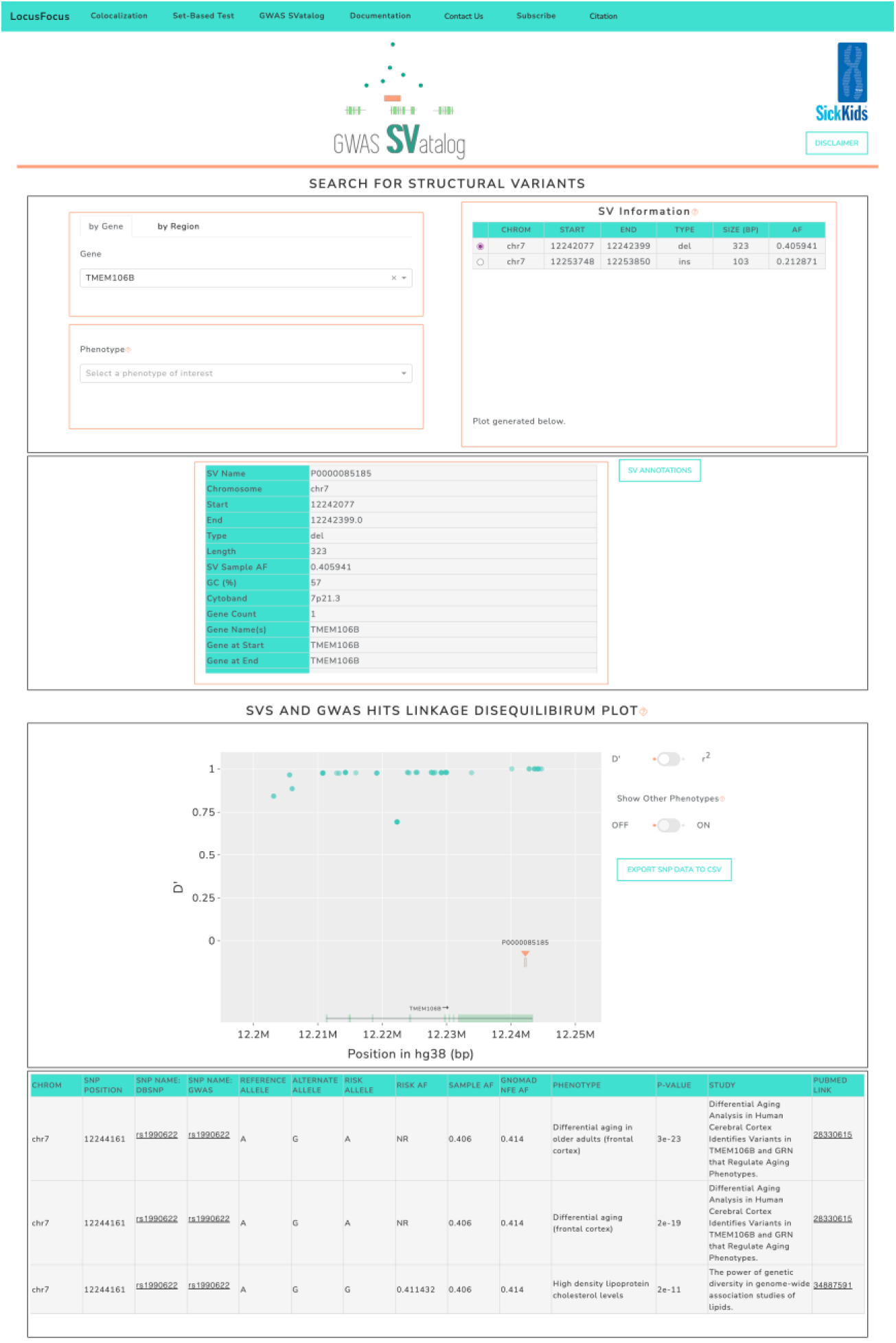
The GWAS SVatalog interface. The navigation bar provides extra information for usage, citation and the *LocusFocus* suite of software tools. The top section of the webpage provides filters to search for a gene, phenotype and/or SV of interest. The middle section gives additional information on the selected SV. The bottom section displays the plot generated based on the filters selected. In addition, if a SNP in the plot is also selected, additional information about this SNP will be populated in a table below the plot.

## Results

### SV calls

We devised a pipeline (Figure 1) that merges SV calls made by pbsv and Sniffles from our PacBio CLR long-read sequencing data, with SV calls from Long Ranger, CNVnator, ERDs, and Manta from our 10XG linked-read sequencing data. In total, 129,485 distinct SVs >50 bp were identified with an average size of 977 bp. The N50 read length range of 22 – 35 kb from the long-reads resulted in most SVs detected within a single read. We observe 60,591 insertions and 63,301 deletions, which is higher than previously reported genome-wide long-read SV call sets (Audano *et al*., 2019; Beyter *et al*., 2021).

From the total 129,485 SV calls, 87,183 distinct SVs were found in at least three individuals from our sample cohort. Each of these SVs have been annotated with their basic information such as length, frequency and type, in addition to more detailed annotations such as percentage of GC content, genes that overlap, repeat region overlap, and percentage overlap with gnomAD populations.

### Generalizability and validation of SV calls

Our SVs were called from the DNA of 101 individuals with CF who are predominately of European origin (Eckford *et al*., 2019) however, these SV calls and frequencies are generalizable to non-CF individuals of European origin. This is because, although the SVs were called from individuals with two CF-causing mutations in the CF transmembrane conductance regulator (*CFTR*) on chromosome 7, we show the genetic background of our cohort does not differ significantly from that of a healthy European population, with the exception of the LD block encompassing *CFTR*.

For this analysis, we compared the genotypes of high-confidence SNPs from the CF cohort to the EUR population of 1000 Genomes Phase 3 (1000 Genomes Project Consortium *et al*., 2015) (Supplementary Figure 1) demonstrating no significant differences in AFs outside of the *CFTR* locus using Fisher’s exact test. In our cohort of 101 individuals, 91 are of European origin while the remaining 10 are from varying demographics including African and Asian. The estimation of ancestry of the cohort was conducted using GrafPop (Jin et al. 2019). The most common CF-causing mutation is F508del, with 51 homozygous carriers and 20 heterozygotes. Additional cohort demographics are provided in Supplementary Table 1.

We compared our SV calls to three publicly available SV sources: HG002 (Zook *et al*., 2016); a 15 multi-ethnic sample cohort using PacBio long-reads (Audano *et al*., 2019); and SVs called in 3,622 Icelandic individuals using Oxford Nanopore long-reads (Beyter *et al*., 2021). The comparison with our SV database was carried out using Truvari based on the NAF of the polymorphism. NAF is the percentage of alleles that deviate from the reference at this SV site. As allelic differentiation has not been made in this dataset, any deviation from the reference genome, GRCh38, in the stated boundary classifies as an SV. For example, an SV in our cohort with an NAF of 0.57 indicates that 57% of the haplotypes in our cohort deviate from GRCh38 at the location of this SV. Of the 8,902 SVs in our population with NAF > 0.5, 94% of these SVs were also called in the three publicly available SV sources we investigated. We then looked at 31,087 SVs with NAF > 0.1 in our population and found 85% of these SVs were also seen in the three comparison sources.

To compare SV genotype calls, we used publicly available HG002 sequences from both 10XG and PacBio CLR, and the HG002 draft benchmark file from GIAB. After running the sequences through our SV calling pipeline, we found an 82% concordance rate using Truvari bench.

When comparing our SV catalog to gnomAD v4.1, a recently updated public database created using short-read sequences, we see a 53.46% overlap. Here we find that a lower percentage of our calls match the SVs in this database (Supplementary Table 2). Similar to the long-read dataset comparison, we subsetted SVs based on the NAF of the polymorphism. From the SVs in gnomAD we identify 68% and 72%, respectively for SVs in our cohort with NAF > 0.1 and NAF > 0.5. This aligns with the assumption that short-read data is limited in the SVs it can genotype. Additionally, from the overlap of SVs, we compared NAF values by calculating the CCC between the two datasets, resulting in 0.736 (95% CI: 0.720 - 0.749).

### Distribution of SVs across the genome

The distribution of SV lengths shows prominent peaks at the 300 bp and 6,000 bp lengths corresponding to *Alu* and LINE elements, respectively (Collins *et al*., 2020) (Supplementary Figure 3). A high density of SV calls can be seen in telomeric regions (Supplementary Figure 4) with most of the singleton calls detected there. The number of SVs overlapping regulatory features, CpG islands, repeats, segmental duplications, and topologically associated domain (TAD) blocks can be found in Supplementary Table 3-4. We see that enhancers are frequently overlapping with SVs. This is intriguing as enhancers affect gene expression within a given TAD block (Panigrahi and O’Malley, 2021) and the TAD block boundaries demarcate the 3D conformation of a locus (McArthur and Capra, 2021). There are 73,655 unique common SVs that are present within TAD blocks, and we observed only 72 SVs to overlap TAD boundaries, which can have functional consequences when altered such as disruption of gene expression (Panigrahi and O’Malley, 2021).

Supplementary Table 4 provides the number of unique SVs that overlap with gene boundaries (i.e. start and/or stop codons) potentially affecting transcription of the gene. The table also includes 1,062 unique SVs overlapping with exon/intron boundaries with the potential of interrupting splice sites and creating non-functional transcripts. We have also noted a small percentage of SVs overlapping entire genes as shown in Supplementary Table 5. As seen in the table, there is one deletion SV encompassing 21 genes. This SV is depicted as a common SV in other publicly available databases such as dbVar (Lappalainen *et al*., 2013) and DGV (MacDonald *et al*., 2014). The *DUB/USP17* gene family is among the 21 genes deleted by this SV. They are a highly conserved family of genes within and among mammalian species consisting of a high proportion of tandem repeats (Burrows *et al*., 2005; Yang *et al*., 2021).

Association testing using logistic regression (Methods detailed in Supplementary materials) indicated that higher SV NAF was associated with lower odds of CpG Island overlap (p<0.0001) and lower odds of Promoter overlap (p=0.0123). SV Type was significantly associated with regulatory feature overlap, although the most/least likely SV types to overlap regulatory features varied between regulatory features (p ranging from <0.0001 to 0.0377). Higher SV size and greater numbers of GWAS SNPs near an SV led to higher odds of SV overlap with each regulatory feature (all p<0.01). SVs that were less directly tagged by GWAS SNPs (based on either D’ or r^2^) tended to overlap CpG islands (p<0.0001) and promoters (all p≤0.0445). A summary of SV length, type and NAF is provided in Supplementary Table 6.

### Identifying GWAS loci that may be explained by an SV using GWAS SVatalog

GWAS SVatalog is capable of queries based on a phenotype of interest or a genomic region of interest in GRCh38 coordinates. Figure 3 provides examples of analyzing a single target SV. Figure 3A provides sample output of LD between a chosen target SV and GWAS-significant SNPs for all phenotypes while Figure 3B shows an alternate view of the target SV, highlighting the LD with GWAS-significant SNPs of a specified target phenotype. To conduct further investigation, all data in the plot, including phenotype associations, can be downloaded by the user.

**Figure 3:**
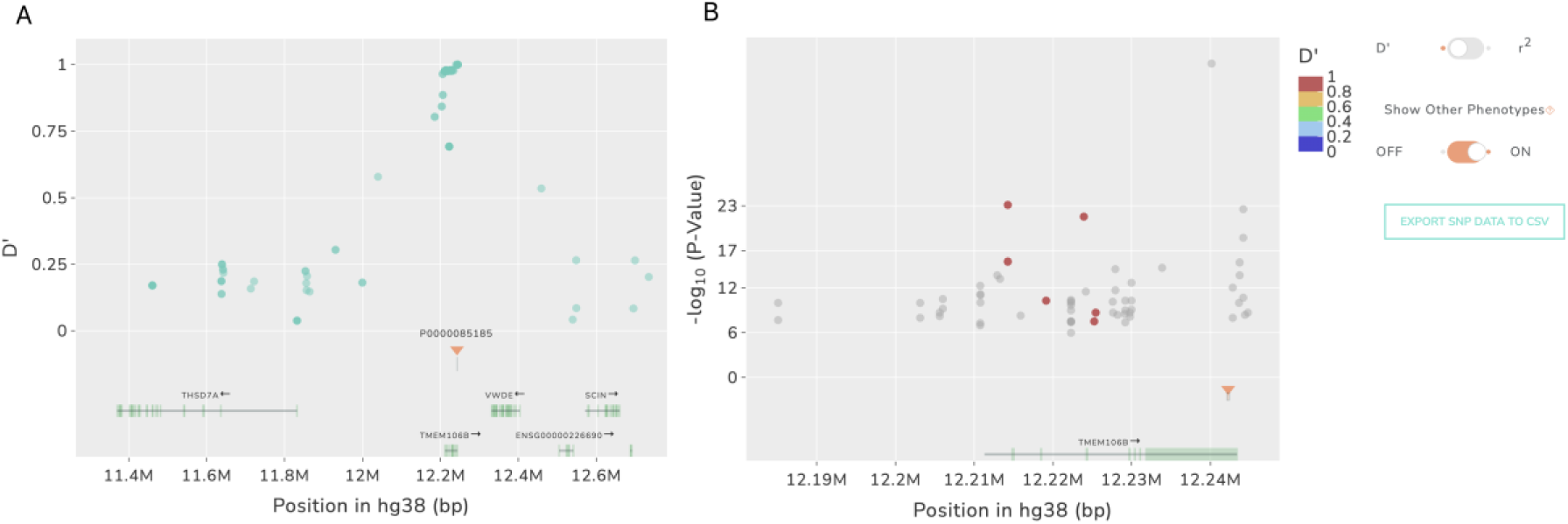
The available displays in GWAS SVatalog to analyze a locus of interest. A) Plot created by only selecting an SV (chr7: 12242077-12242399) of interest (no phenotype selected). The blue dots are SNPs from the GWAS Catalog, and the y-axis is the LD relationship (D’ or r^2^) to the selected SV. B) Plot created by selecting the same SV of interest along with a specific phenotype (depression). The dots are colored by their LD relationship (D’ or r^2^) to the selected SV and GWAS-significant SNPs of the selected phenotype. SNPs can appear more than once when different studies result in varying p-values for the significance of the SNP. The y-axis is based on the –log_10_(p-value) provided by the studies in the GWAS Catalog. The gray dots show GWAS-significant SNPs of other phenotypes for the selected SV.

An overview of the distribution of SVs available in GWAS SVatalog based on their type (deletion, duplication, insertion and inversion) and annotation by functional genetic regions can be found in Table 1. Supplementary Table 7 provides the proportion of SVs associated with GAL using the max LD score per SV. There are 9,438 SVs, located within a gene, in high LD (D’ ≥0.8) with GAL, with 530 GWAS-significant SNPs being exonic. Of the 36,295 SVs with MAF ≥ 0.1, 35,732 SVs had SNPs within 1Mb of their boundaries with non-zero LD. In total, there are 116,870 GAL. Among these, 64,919 GAL are in high LD (D’ ≥ 0.9) with 21,578 SVs.

**Table 1:**
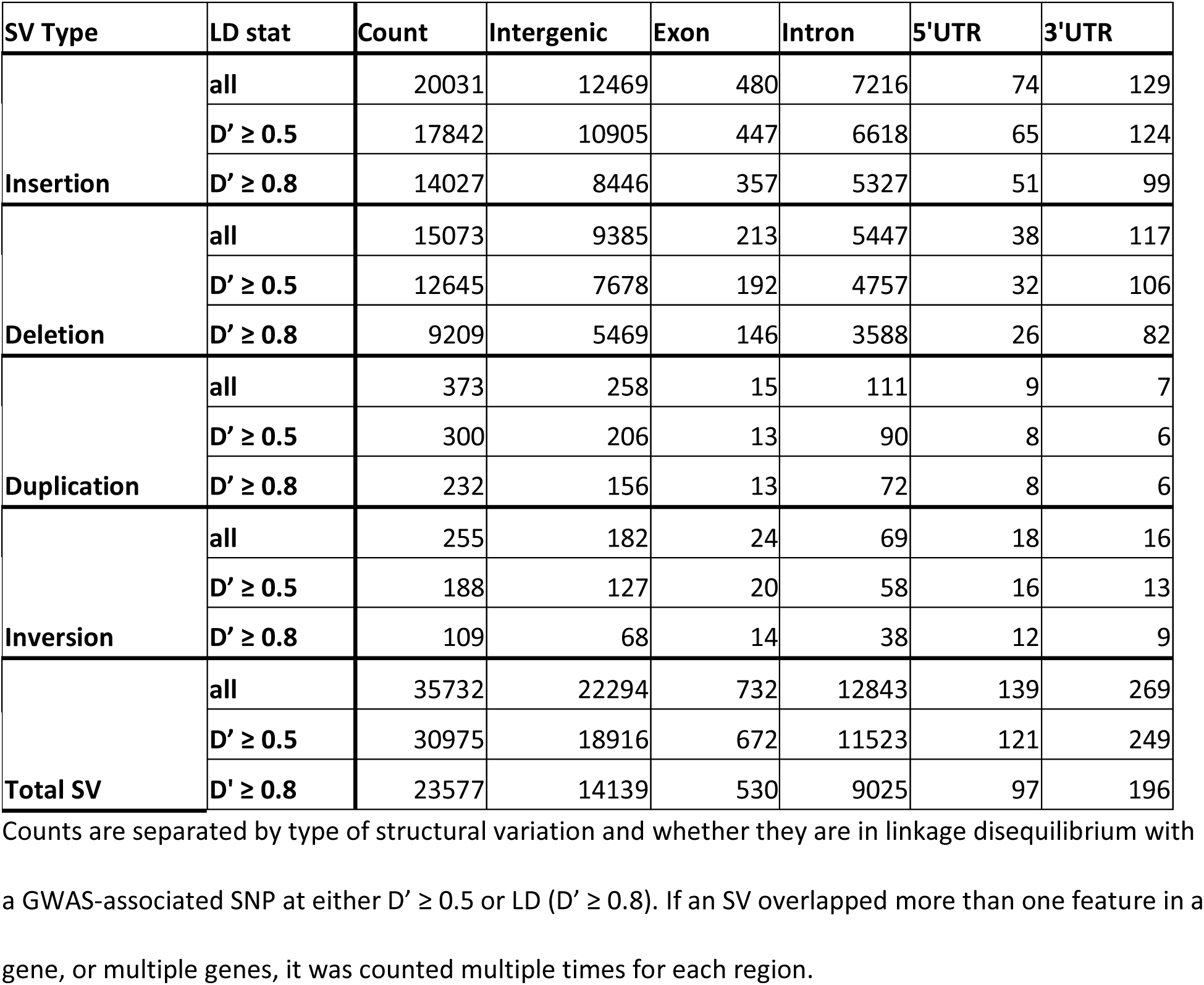
Distribution of structural variants across genomic regions.

Using the GWAS SVatalog and the SV calls that we generated, we wanted to identify SVs that may explain GAL. We first filtered 21,578 SVs in high LD (D’ ≥ 0.9) with GAL from where we only considered 12,002 SVs that overlap protein-coding genes. Of these, we focused on 9,914 SVs in high LD with two or more phenotypes (i.e. pleiotropic/replication). We then randomly selected 100 SVs from this subset and visualized each SV using GWAS SVatalog. We identified the previously reported causal deletion SV present in the 3’UTR of *ARMS2* associated with age-related macular degeneration (Fritsche *et al*., 2008; Micklisch *et al*., 2017) (Figure 4A). We also identified three novel candidates’, shown in Table 2 and Figure 4B-D, where an SV appears to potentially affect regulatory activity which could explain the GWAS signals.

**Figure 4:**
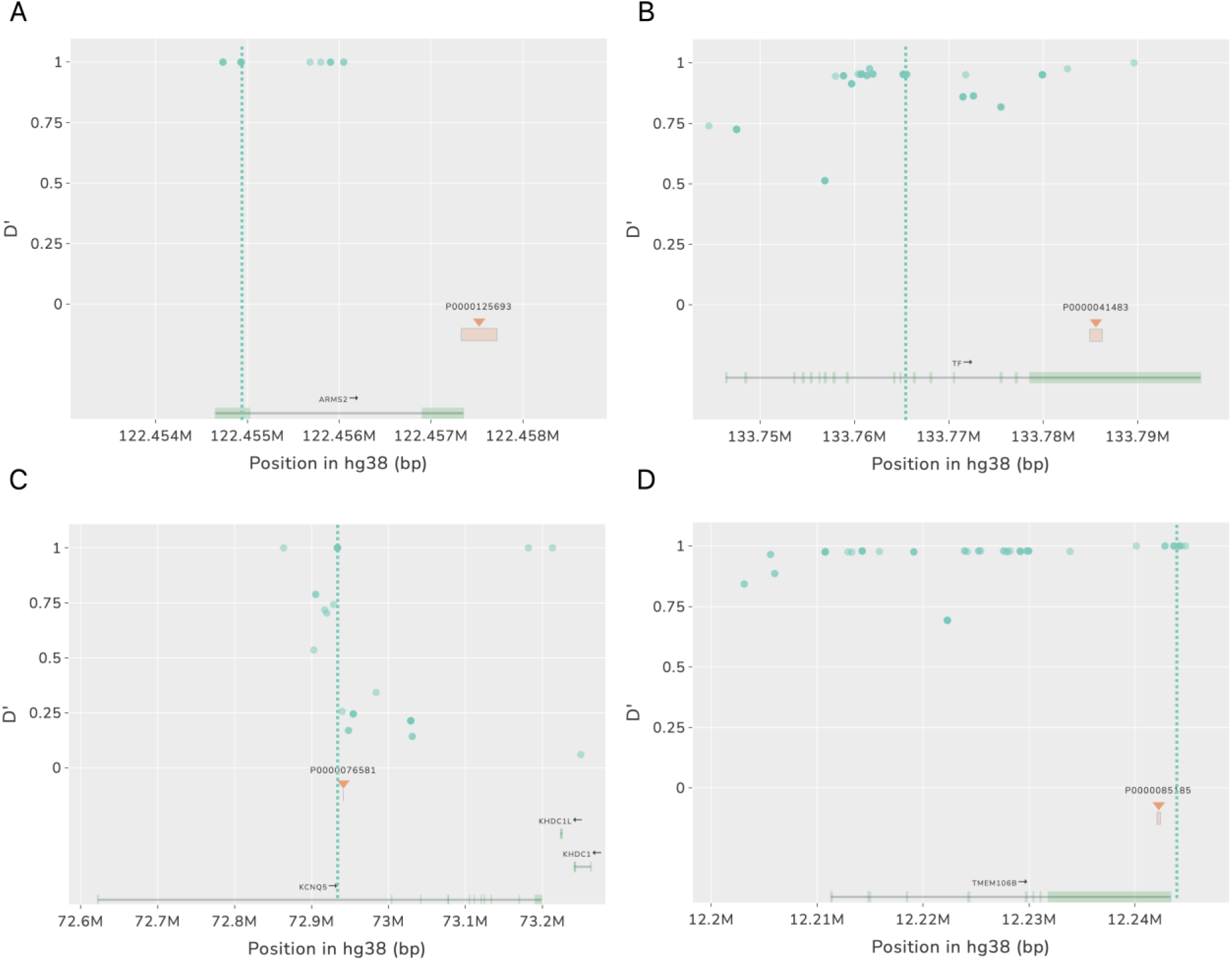
Screenshots of plots created by GWAS SVatalog to analyze GAL. The dotted line indicates the location of the SNP previously shown to be associated in this region for specific phenotypes. A) is an example of a causal SV previously identified in high LD with a GWAS-significant SNP while B) - D) are putative examples of SVs found using GWAS SVatalog that could potentially impact functionality in the GAL. A) rs10490924 associated in GWAS for age-related macular degeneration (AMD). The SV, in high LD, located in the 3’UTR of ARMS2 has been shown to be a causal factor for AMD. B) A 1317 bp SV in high LD with rs3811647, an associated SNP in serum transferrin level GWAS C) A 54 bp SV in high LD with rs7744813, an associated SNP in refractive error GWAS D) A 323 bp deletion in high LD with rs1990622, an associated SNP in Alzheimer’s disease.

**Table 2:** Three candidate causal SVs identified by fine-mapping using GWAS SVatalog.

|  |  | Example 1 | Example 2 | Example 3 |
| --- | --- | --- | --- | --- |
| **Gene** |  | *TF* | *KCNQ5* | *TMEM106B* |
| **Phenotype** |  | serum transferrin levels | myopia and refractive error | multiple neurodegenerative diseases |
| **SV** | **Chromosome Location** | 3:133784922-133786238 | 6:72941318-72941371 | 7:12242077-12242399 |
|  | **Genomic Location** | 3'UTR | intron | 3'UTR |
|  | **Type** | deletion | deletion | deletion |
|  | **Length (bp)** | 1317 | 54 | 323 |
|  | **AF** | 0.342 | 0.114 | 0.406 |
|  | **Feature Overlap** | transcription factor *SREBP2* binding site, SINE-VNTR-*Alu* retrotransposon | H3K4me1 and H3K27ac, *ZNF263* and *PCBP2* transcription factor ChIP-seq clusters | SINE, H3K4me1, H3K4me3, H3K27ac, CpG island, *TBX21* and *PKNOX1* transcription factor binding sites |
| **SNP** | **Name** | rs3811647 | rs7744813 | rs1990622 |
|  | **Chromosome Location** | 3:133765185 | 6:72933566 | 7:12244161 |
|  | **Genomic Location** | intron | intron | downstream of gene |
|  | **AF** | 0.366 | 0.614 | 0.406 |
| **LD Statistics** | **D'** | 0.952 | 1 | 1 |
|  | **r<sup>2</sup>** | 0.814 | 0.081 | 1 |

The first candidate SV (Figure 4B), a 1317 bp deletion (NAF = 0.342) localized at the 3’UTR of *TF* on chromosome 3 was found to be in LD (D’ = 0.952) with rs3811647 (AF=0. 366), a GWAS-significant SNP in an iron biomarker GWAS (Benyamin *et al*., 2009). This SV is a SINE-VNTR-*Alu* (SVA) retrotransposon that removes a *SREBP2* binding site. The second candidate (Figure 4C), a 54 bp deletion (NAF = 0.114) in intron 1 of *KCNQ5* on chromosome 6 was in high LD (D’=1) with rs7744813 (AF = 0.614), associated with refractive error (Verhoeven *et al*., 2013). This intronic SV has the potential to influence *KCNQ5* expression as it is located within H3K4me1 and H3K27ac histone marks and two transcription factor ChIP-seq clusters (*ZNF263* and *PCBP2*) known to be transcriptional and translational suppressors, respectively (Ren *et al*., 2016; Yu *et al*., 2020). The third candidate (Figure 4D), a 323 bp deletion localized to the 3’UTR of *TMEM106B* on chr7 (NAF = 0.406) is in perfect LD (D’ = 1) with rs1990622 (AF = 0.406), associated with Alzheimer’s disease (Hu *et al*., 2021). This SV is a SINE element that removes multiple epigenetic elements: three histone marks (H3K4Me1, H3K4Me3, and H3K27Ac), a CpG island, and two transcription factor binding sites (*TBX21* and *PKNOX1*).

In these examples, the GAL have no known functional consequence tying phenotype to genotype, to the best of our knowledge. GWAS SVatalog provided an alternative explanation for these GAL by visualizing SVs in high LD that could be putatively functional. These are but a small subset of examples identified from a large list of candidate SVs. Further investigation using GWAS SVatalog has the potential to identify many other SVs that could explain GALs and lead to translational discoveries.

## Discussion

Here we introduce a novel web tool, GWAS SVatalog, that integrates SVs into visualizations of GAL to assist in fine-mapping and putative causal variation identification. We created a catalog of SVs from a cohort of individuals representative of populations of European origin using a combination of PacBio CLR sequencing and 10XG linked-read sequencing. By utilizing both of these technologies, we are able to leverage their best attributes: CLR can identify large polymorphisms, like SVs, and 10XG excels in genotyping SNPs. The SV call set we produced, and corresponding LD calculations with respect to SNPs reported in the GWAS Catalog, has been made available through the GWAS SVatalog visualization tool. The web tool aids fine-mapping by incorporating 35,732 SVs at GAL and visualizing the LD relationship between these SVs and 116,870 GWAS-significant SNPs from 14,479 different human traits.

For our cohort, we demonstrated that the AFs across the genome (with the exception of the *CFTR* locus) are representative of the EUR sub-group of the 1000 Genomes healthy control group (Supplementary Figure 1). This gives us confidence in the ability of GWAS SVatalog to represent accurate LD relationships for a GWAS comprised of a high proportion of Europeans. For other ethnicities, this remains a major limitation of the tool which we are hoping to address by including more diverse SV call sets as they are generated, such as those included in the All of Us study (All of Us Research Program Genomics Investigators, 2024).

In 2020, gnomAD released a dataset linking common SVs genotyped from short-read sequences with GWAS variants (Collins *et al*., 2020). However, SV identification is limited when genotyped using short-read sequence data due to their average read length of 150 bps preventing larger SVs that can span thousands of base pairs to be called efficiently (Mahmoud *et al*., 2019). Short-reads also have difficulty calling SVs that lie in repeat regions (Collins *et al*., 2020; Kosugi and Terao, 2024), are related to copy number variations (Collins *et al*., 2020), or have a combination of events (complex) such as inversion-duplication (Sedlazeck *et al*., 2018; Collins *et al*., 2020). Overall, short-read sequences have been shown to miss about 30% of SV calls compared to long-read sequences (Sedlazeck *et al*., 2018). Supplementary Table 2 shows that our data aligns with this estimate as well when comparing the newest short-read gnomAD SV database to our long-read SV database.

Benchmarking conducted on our SV database against three publicly available long-read SV resources (Zook *et al*., 2016; Audano *et al*., 2019; Beyter *et al*., 2021) supports the reliability of our SV calls as 85% of the common SVs (NAF > 0.1) were found in the other three datasets. The three sources were derived from varying sequencing platforms and software calling methods, contributing to the differences in the SVs identified.

Association testing showed that SV NAF was inversely associated with overlap of CpG islands and promoters, which is consistent with existing literature (Schloissnig et al. 2025). Both CpG island overlap and promoter overlap were also associated with SV-SNP LD. Specifically, SVs that are less directly tagged by GWAS SNPs (whether measured by maximum D’ or r^2^) were more likely to overlap CpG islands or promoters than to overlap no regulatory feature, after controlling for SV size, type, and NAF. This is consistent with the hypothesis that some GWAS hits may be driven by SVs altering the function of nearby genes.

We used GWAS SVatalog to identify three candidate SVs that may explain the GWAS signal reported in the literature, since the functional polymorphism was not identified to our knowledge. These SVs lie in regions within genes and/or are overlapping regulatory features that could affect the level of gene expression. For example, the 54 bp deletion SV in the first intron of *KCNQ5*, is in high LD (D’ = 1) with an intronic variant, rs7744813. This SNP has not been shown to be a causal variant but is repeatedly shown to be significantly associated in GWASs of refractive error and myopia (Verhoeven et al. 2013; Li et al. 2021; Liao et al. 2017). As this SV removes part of two transcription factor binding sites, *ZNF263* and *PCBP2,* which have previously been shown to be transcriptional and translational suppressors (Ren *et al*., 2016; Yu *et al*., 2020), further analysis of their interaction with *KCNQ5* could potentially demonstrate the significance of the GWAS signal at this locus. The second candidate SV identified using GWAS SVatalog is a 323 bp deletion in the 3’UTR of the gene *TMEM106B*. This SV is in complete LD (D’ = 1 and r^2^ = 1) with a variant, rs1990622, located downstream of the gene. Even though rs1990622 has been shown to be highly associated in GWAS of Alzheimer’s disease (Hu *et al*., 2021), it is also known to impact other neurological disorders such as frontotemporal dementia and Parkinson’s disease (Van Deerlin *et al*., 2010; Lee *et al*., 2023). Studies have linked rs1990622-A to an increase in levels of TMEM106B protein in the lysosome leading to an increased risk of dementia (Lee *et al*., 2023). As this SNP and the SV are in complete LD, functional studies of this SV could determine whether the SV could affect the neurological phenotypes. For the third candidate SV, a 1317 bp deletion is seen in high LD (D’ > 0.95) with an intronic variant rs3811647 that has been repeatedly associated with a GWAS of variation of serum transferrin levels (Benyamin *et al*., 2009; Pichler *et al*., 2011; Blanco-Rojo *et al*., 2012). Transferrin is essential for the homeostasis of iron levels by aiding in its transportation to cells (Blanco-Rojo *et al*., 2012) and the role of rs3811647 in the expression of the transferrin gene (*TF)* has not yet been reported. As this SV is a SINE-VNTR-*Alu* (SVA) retrotransposon and encompasses a few transcription factor binding sites (including *SREBP2*), further functional analyses will determine if this SV impacts *TF* expression and variation of serum transferrin. As supporting evidence, SVAs have been reported to have a pronounced effect on gene expression (Hancks and Kazazian, 2010; Quinn and Bubb, 2014) and *SREBP2* has been shown to directly affect the transcription of *TF* in circulating tumor cells (Hong *et al*., 2021). Therefore, upon functional follow-up, this SV could explain the significance of the GWAS signal. All three candidate SV examples illustrate how GWAS SVatalog can improve fine-mapping of a locus without genotyping of the SVs in the original GWAS discovery sample and can direct future studies for causal variant identification.

Although GWAS SVatalog provides a unique contribution to fine-mapping moving forward, we note several limitations. First, the data produced for GWAS SVatalog uses SV calls from reference-aligned long-reads which improves SV identification in contrast to short reads but is still dependent on alignments that are susceptible to reference bias. Reference bias is a systematic error that occurs during the alignment of sequencing reads to a reference genome resulting in underrepresentation of variation that differs significantly from the reference genome. It produces errors such as false-negatives, false-positives, and incomplete detection of complex SVs (Günther and Nettelblad, 2019; Martiniano *et al*., 2020; Valiente-Mullor *et al*., 2021). More specifically, the development of the GWAS SVatalog tool was motivated by a 20 kb deletion polymorphism tagged by a suggestive SNP in a GWAS of intestinal obstruction in CF (Gong *et al*., 2019; Mastromatteo *et al*., 2023). This finding led us to hypothesize that similar cases might exist in other contexts, where GWAS-significant SNPs are not the variation that explain the GWAS signal but rather markers of larger, functionally significant variations. Despite being the motivation of this work, we were unable to accurately call the 20 kb polymorphism using PacBio CLR data aligned to GRCh38. Even with PacBio HiFi data (data not shown here), we were still unable to call the 20 kb insertion against GRCh38 despite the increase in base pair accuracy. Therefore, the SVs cataloged here are an incomplete subset of all SVs in our population. Future work with *de novo* assemblies and reference pangenomes can potentially improve SV calling by mitigating the effect of reference bias.

The second limitation arises from the SV calling software currently available. We were unable to fully characterize the alleles of each SV due to difficulty in calling their boundaries. Due to this challenge, we collapsed all SVs in the same position across samples into one non-reference SV allele, leading us to treat all SVs in this dataset as biallelic. As a result of collapsing non-reference alleles together, the reported NAF of the SVs does not fully capture the nuance of each allele. As some SVs will be multiallelic, we are unable to showcase that variation for now. However, even if the SVs were captured to be multiallelic, there is no standard methodology to calculate LD for a pair of loci belonging to biallelic and multiallelic variants, bringing us to the third limitation of all polymorphisms in this dataset being treated as biallelic. Due to this, the resulting LD between GWAS-significant SNPs and SVs may be slightly overestimated (Hedrick, 1987; Gaunt *et al*., 2006; Zhao *et al*., 2007; Jiang *et al*., 2020). However, the D’ coefficient is a reliable statistic with respect to the allelic discrepancy as its observation has been shown to be predominately frequency independent (Hedrick, 1987; Zapata, 2000; Zhao *et al*., 2007). As a significant proportion of SV-SNP pairs have low r^2^ and high D’, SNPs seem to be tagging SVs with differing frequencies from SNPs, possibly reflecting the multi-allelic nature of SVs.

The integration of SVs in GAL will fill the gap in knowledge created by primarily using SNPs in GWAS. Using GWAS SVatalog to guide functional follow-up by visualizing LD between common SVs and GALs will aid in fine-mapping to better understand the mechanism of action at the GAL. We are confident in the curation of our SV database built primarily from long-reads and hope to make future improvements by using HiFi sequences and pangenome references to alleviate the challenges imposed by SV genotyping and reference bias. In the future we hope to expand the long-read sequencing data used to call the SVs that are cataloged in GWAS SVatalog, to ensure a more comprehensive SV catalog that can inform GAL in diverse populations.

## Acknowledgements

We thank the patients, care providers and clinic research assistants, collaborators, and principal investigators involved in CF Centers throughout Canada for their contributions to the CF Canada Patient Registry and CF Canada-Sick Kids Program in Individual Therapy. The authors wish to acknowledge the staff supporting the High Performance Computing cluster, Research Helpdesk department and The Centre for Applied Genomics at The Hospital for Sick Children, Toronto.

## Authors’ contributions

Conceptualization: L.J.S. Sample recruitment: K.K., J.A., P.E., F.R. Sample processing: F.L., K.K. Data processing: S.C., Z.W., S.M., B.T., T.M., G.P., R.P., W.S., S.A., J.W. Formal analysis: S.C., Z.W. Funding acquisition: L.J.S. Investigation: S.C., Z.W., B.T., S.M., E.S., L.J.S. Methodology: S.C., Z.W. Project administration: L.J.S. Supervision: L.J.S. Visualization: S.C. Writing-original draft: S.C., Z.W., L.J.S. Writing, review, and editing: all authors.

## Data archiving

The CFIT sequencing data used in this study is described in Eckford *et al*. 2019 and is available through the Canadian CF registry at https://www.cysticfibrosis.ca/our-programs/cf-registry/requesting-canadian-cf-registry-data. The datasets generated and/or analyzed during the current study are available in the Zenodo repository https://doi.org/10.5281/zenodo.13367574. GWAS SVatalog is available at https://svatalog.research.sickkids.ca/. The source code can be found at https://github.com/strug-hub/gwas-svatalog. The documentation for the tool is available at https://gwas-svatalog-docs.readthedocs.io/en/latest/index.html.

## Research ethics statement

SickKids Program in Individualized CF Therapy (CFIT) was approved by the Research Ethics Board of the Hospital for Sick Children (#1000044783 from 2014 to present) and sub-site St. Michael’s Hospital (#14 188 from 2015 to present). Written informed consent was obtained from all participants or parents/guardians/substitute decision makers prior to inclusion in the study.

