## Supplementary Table 6 for "GWAS SVatalog: a visualization tool to aid fine-mapping of GWAS loci with structural variations"

**Supplementary Table 6: Summary of each SV's max D', length, type, and NAF.**

| Dbin | SV_Type | N | Mean_Length | Median_Length | Range_Length | Mean_NAF | Median_NAF | Range_NAF | Length_50_100 | Length_100_500 | Length_500_1000 | Length_1000_5000 | Length_5000_max | NAF_0_0.2 | NAF_0.2_0.4 | NAF_0.4_0.6 | NAF_0.6_0.8 | NAF_0.8_1 |
|---|---|---|---|---|---|---|---|---|---|---|---|---|---|---|---|---|---|---|
| 0-0.2 | Deletion | 167 | 1,247 | 134 | 50-86,993 | 0.358 | 0.332 | 0.109-0.866 | 39.52% | 39.52% | 5.99% | 11.38% | 3.59% | 20.36% | 43.11% | 24.55% | 10.18% | 1.80% |
| 0-0.2 | Duplication | 9 | 3,324 | 1,735 | 52-11,989 | 0.332 | 0.347 | 0.134-0.703 | 11.11% | 0.00% | 0.00% | 66.67% | 22.22% | 33.33% | 44.44% | 11.11% | 11.11% | 0.00% |
| 0-0.2 | Insertion | 222 | 553 | 260 | 101-16,137 | 0.415 | 0.413 | 0.104-0.886 | 0.00% | 74.32% | 15.77% | 8.56% | 1.35% | 18.47% | 29.28% | 33.78% | 13.96% | 4.50% |
| 0-0.2 | Inversion | 13 | 74,057 | 903 | 390-933,266 | 0.376 | 0.307 | 0.124-0.881 | 0.00% | 7.69% | 46.15% | 15.38% | 30.77% | 30.77% | 30.77% | 23.08% | 7.69% | 7.69% |
| 0.2-0.4 | Deletion | 1035 | 716 | 130 | 50-102,603 | 0.399 | 0.381 | 0.104-0.896 | 40.39% | 44.15% | 5.51% | 7.15% | 2.80% | 13.72% | 39.42% | 32.46% | 12.56% | 1.84% |
| 0.2-0.4 | Duplication | 23 | 2,320 | 405 | 51-22,255 | 0.373 | 0.322 | 0.104-0.807 | 21.74% | 30.43% | 17.39% | 21.74% | 8.70% | 17.39% | 43.48% | 21.74% | 13.04% | 4.35% |
| 0.2-0.4 | Insertion | 986 | 357 | 238 | 101-4,458 | 0.422 | 0.406 | 0.104-0.886 | 0.00% | 81.14% | 14.20% | 4.67% | 0.00% | 11.16% | 36.92% | 35.19% | 14.81% | 1.93% |
| 0.2-0.4 | Inversion | 35 | 23,790 | 533 | 147-755,238 | 0.347 | 0.337 | 0.104-0.752 | 0.00% | 48.57% | 8.57% | 25.71% | 17.14% | 25.71% | 40.00% | 22.86% | 11.43% | 0.00% |
| 0.4-0.6 | Deletion | 1967 | 865 | 119 | 50-155,155 | 0.385 | 0.356 | 0.104-0.896 | 43.72% | 42.40% | 5.69% | 6.25% | 1.93% | 17.69% | 39.60% | 28.22% | 12.91% | 1.58% |
| 0.4-0.6 | Duplication | 38 | 2,164 | 506 | 51-29,810 | 0.304 | 0.245 | 0.104-0.797 | 21.05% | 26.32% | 21.05% | 21.05% | 10.53% | 36.84% | 39.47% | 15.79% | 7.89% | 0.00% |
| 0.4-0.6 | Insertion | 2165 | 372 | 247 | 101-8,596 | 0.418 | 0.401 | 0.104-0.896 | 0.00% | 78.24% | 16.91% | 4.62% | 0.23% | 11.73% | 37.92% | 33.16% | 15.57% | 1.62% |
| 0.4-0.6 | Inversion | 43 | 14,485 | 2,107 | 79-179,950 | 0.343 | 0.312 | 0.104-0.748 | 4.65% | 27.91% | 9.30% | 18.60% | 39.53% | 30.23% | 34.88% | 25.58% | 9.30% | 0.00% |
| 0.6-0.8 | Deletion | 1990 | 575 | 124 | 50-111,726 | 0.366 | 0.319 | 0.104-0.896 | 41.71% | 44.27% | 5.48% | 6.48% | 2.06% | 24.97% | 37.14% | 22.36% | 12.81% | 2.71% |
| 0.6-0.8 | Duplication | 20 | 801 | 724 | 51-2,573 | 0.323 | 0.248 | 0.124-0.787 | 20.00% | 20.00% | 20.00% | 40.00% | 0.00% | 35.00% | 35.00% | 15.00% | 15.00% | 0.00% |
| 0.6-0.8 | Insertion | 2631 | 363 | 244 | 101-5,144 | 0.408 | 0.391 | 0.104-0.896 | 0.00% | 79.86% | 15.58% | 4.52% | 0.04% | 12.24% | 38.69% | 33.41% | 14.14% | 1.52% |
| 0.6-0.8 | Inversion | 55 | 28,408 | 681 | 69-1,097,568 | 0.384 | 0.332 | 0.109-0.866 | 1.82% | 38.18% | 16.36% | 21.82% | 21.82% | 18.18% | 41.82% | 25.45% | 12.73% | 1.82% |
| 0.8-1 | Deletion | 8353 | 1,015 | 142 | 50-517,542 | 0.302 | 0.223 | 0.104-0.896 | 38.08% | 43.79% | 6.17% | 8.64% | 3.32% | 44.46% | 30.96% | 12.25% | 8.55% | 3.78% |
| 0.8-1 | Duplication | 154 | 4,941 | 460 | 51-316,834 | 0.258 | 0.208 | 0.104-0.891 | 15.58% | 35.71% | 11.04% | 21.43% | 16.23% | 46.75% | 36.36% | 13.64% | 1.95% | 1.30% |
| 0.8-1 | Insertion | 14027 | 422 | 234 | 101-11,805 | 0.34 | 0.267 | 0.104-0.896 | 0.00% | 77.83% | 15.06% | 6.47% | 0.63% | 36.17% | 32.02% | 16.29% | 10.24% | 5.28% |
| 0.8-1 | Inversion | 109 | 21,193 | 1,864 | 78-450,884 | 0.356 | 0.272 | 0.104-0.896 | 2.75% | 28.44% | 9.17% | 24.77% | 34.86% | 36.70% | 32.11% | 11.93% | 11.93% | 7.34% |
