## Supplementary Material for "GWAS SVatalog: a visualization tool to aid fine-mapping of GWAS loci with structural variations"

### SV call frequencies reflect individuals of European origin

Supplementary Figure 1 shows how the variants present in the *CFTR* locus have varying allele frequencies when SNPs from the CF cohort are compared to the EUR population from 1000 Genomes Phase 3. We utilized the list of high-confidence SNPs from Illumina Omni 2.5-8 v1.5 (https://webdata.illumina.com/downloads/productfiles/humanomni25/v1-5/infinium-omni2-5-8v1-5-a1-manifest-file-csv.zip) to ensure the accuracy of most SNPs called from each population, excluding 33 SNPs due to insufficient genotype calling accuracy. Using PLINK1.9, Fisher's exact test was carried out using the genotype data from both populations.

**Supplementary Figure 1: A Manhattan plot comparing allele frequencies of high-confidence SNPs between our CF cohort and the European population of 1000 Genomes Phase 3.** Each dot is a SNP, and the orange dots are SNPs annotated to the CFTR locus (chr7: 117480025-117668665). The grey dotted line marks the level of significance at $-\log_{10}(p) = 8$.

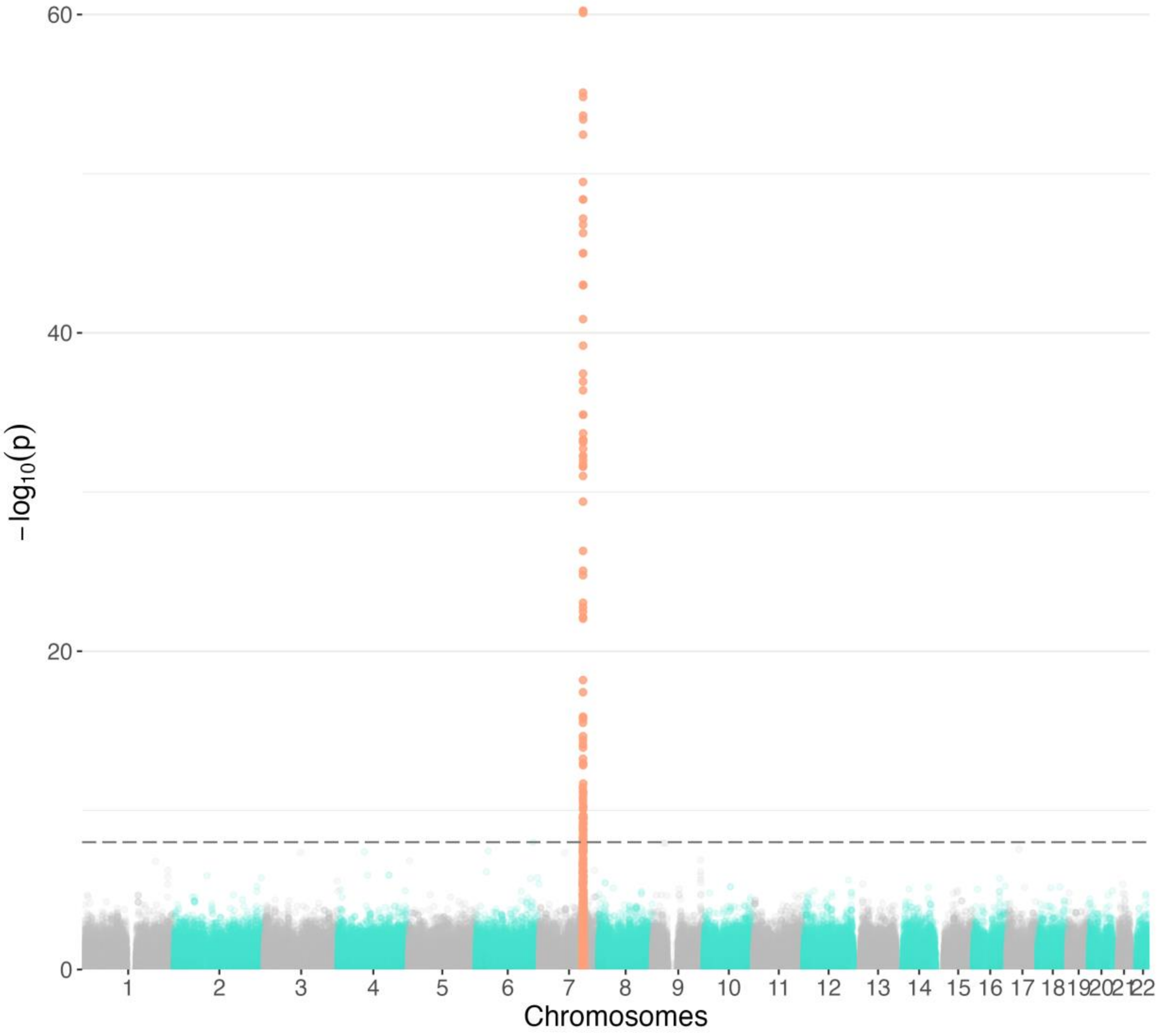

**Supplementary Figure 2: A flowchart showing how data was merged to create GWAS SVatalog.**

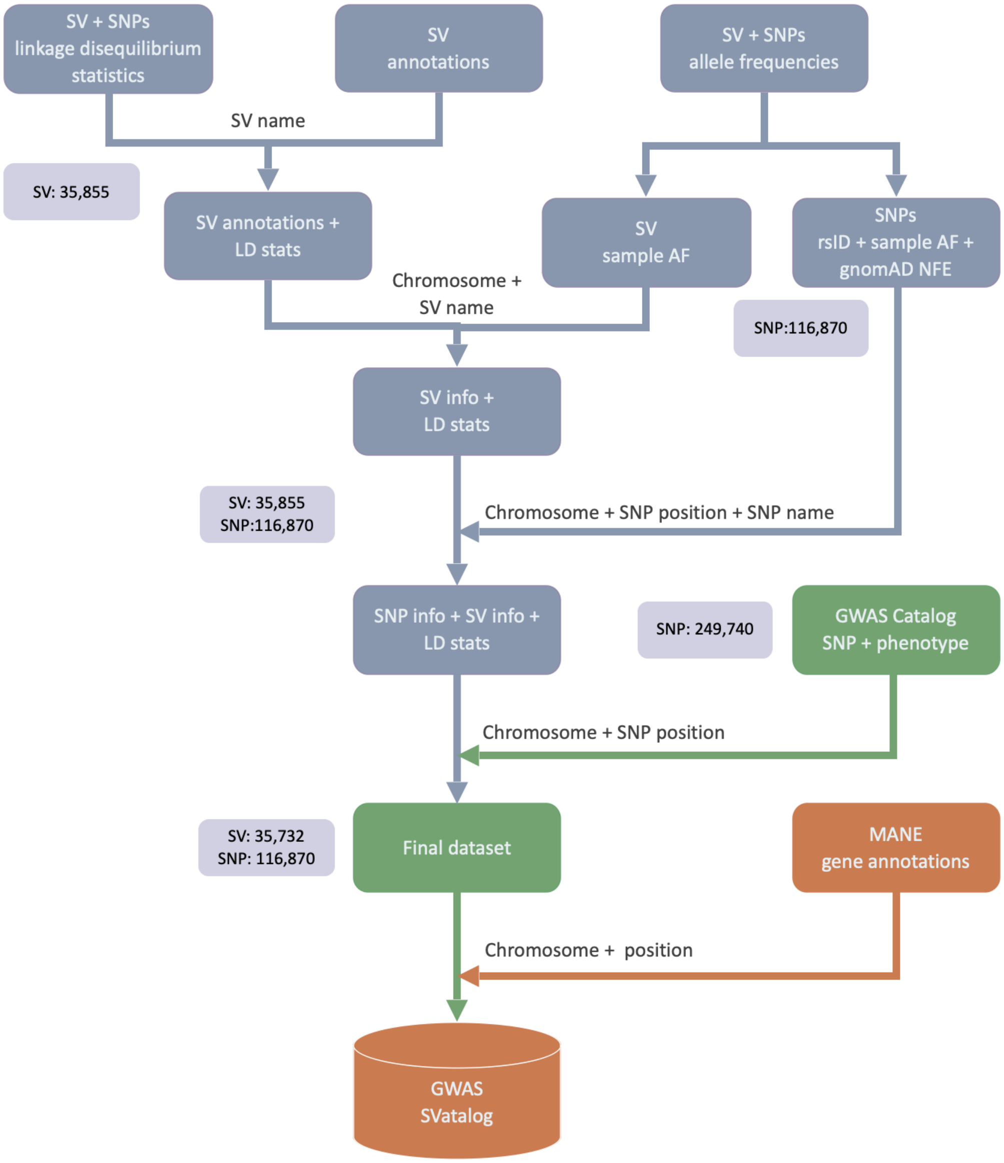

**Supplementary Table 1: Demographic overview of the 101 individuals with CF samples sequenced for this study.**

| | Subcategories | Count |
|---|---|---|
| **Diagnosis Age** | 0 - 10 | 92 |
| | 10 - 20 | 3 |
| | > 20 | 6 |
| **Sex** | Female | 48 |
| | Male | 53 |
| **Canadian Province** | Ontario | 93 |
| | Other | 8 |
| **Ethnicity** | European | 91 |
| | African | 2 |
| | Asian | 2 |
| | Other | 5 |
| | Unspecified | 1 |
| ***CFTR* Genotype** | ΔF508 / ΔF508 | 51 |
| | ΔF508 / Other | 20 |
| | Other / Other | 30 |
| **Pancreatic Insufficient** | Yes | 92 |
| | No | 8 |
| | NA | 1 |

**Supplementary Table 2: Comparison of our SV callset to short-read whole-genome sequence SV calls from gnomAD v4.1.** Total sample size in our SV database is 101 while gnomAD v4.1 is 63,046. The total SVs called in our dataset (including singletons) is 129,485. Total SVs present in gnomAD is ~1.2 million.

| Our SV database | Intersection of SV Calls in gnomAD v4.1 |
|---|---|
| Total SVs ≥ 0.1 NAF | 68.2% |
| Total SVs ≥ 0.5 NAF | 71.6% |
| Singleton calls - European samples | 51.4% |
| Singleton calls - non-European samples | 65.1% |
| Catalogued 87,1183 SVs | 53.5% |

**Supplementary Figure 3: Distribution of SV lengths.** Each line represents the distribution of each SV type, insertion, deletion, duplication, and inversion. The total number of duplication and inversion SVs are significantly lower than insertions and deletions, the distribution appears to be minimal and uniform across lengths. Peaks at 300 bp and 6,000 bp correspond to Alu and LINE elements.

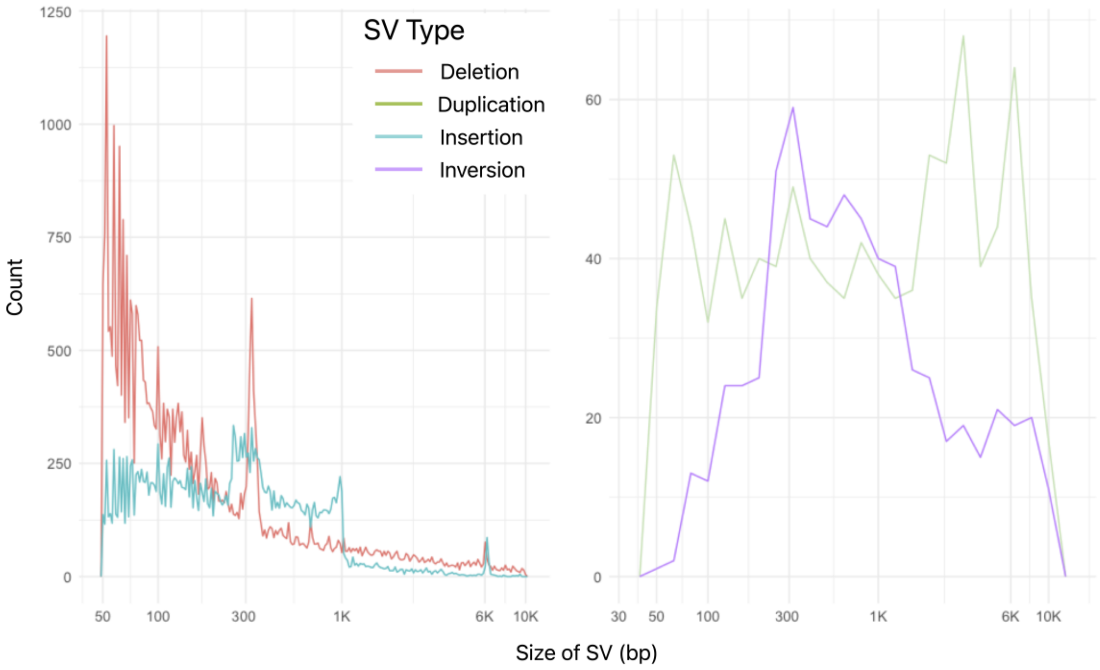


### SV variant calling software comparison

The majority of long-read aligners use a similar seed-chain-extend algorithm (Sahlin *et al.*, 2023). The differences between them lie in the strategies for the alignment itself. Similarly, the range of SV callers available predominantly use a split-read approach to detect SVs while their strategies on deciphering calls result in their differences. Benchmarking a variety of SV calling software has shown that certain tools have higher accuracy for differing complexities of SVs (Kosugi *et al.*, 2019). For example, pbsv ("PacificBiosciences/pbsv: pbsv - PacBio structural variant (SV) calling and analysis tools") focuses on the five simple SVs (insertions, deletions, duplication, inversions, and translocations), Sniffles (Sedlazeck *et al.*, 2018) can detect complex variants such as deletion-insertion combinations, and cuteSV (Jiang *et al.*, 2020) internally merges small indels creating larger structural variations. However, the challenging aspect for each algorithm is determining the boundaries of each SV accurately. The use of different aligners could certainly affect the results of each caller as well.

We used two different combinations of software to achieve a wider range of SV calls on our PacBio CLR sequences; pbmm2 1.1.0 + pbsv 2.2.2 (Pacific Biosciences 2018) and NGMLR 0.2.8 (Sedlazeck *et al.*, 2018) + Sniffles 1.0.11 (Sedlazeck *et al.*, 2018). For our 10XG sequences, we used a wide mix of software: Long Ranger 2.2.2 (10X Genomics 2020), CNVnator (Abyzov *et al.*, 2011), ERDs (Zhu *et al.*, 2012) and Manta (Chen *et al.*, 2016).

**Supplementary Figure 4: The density of the SVs across the genome (autosomes and chrX) identified from 101 individuals with CF are shown in blue.** Each chromosome has been split into one million segments. The density of SVs in each segment is depicted by the shade of blue; the darker the color, the greater the density of SVs. The presence of no SVs in a segment will be shown as white. The orange region above each chromosome corresponds to the location of the centromere.

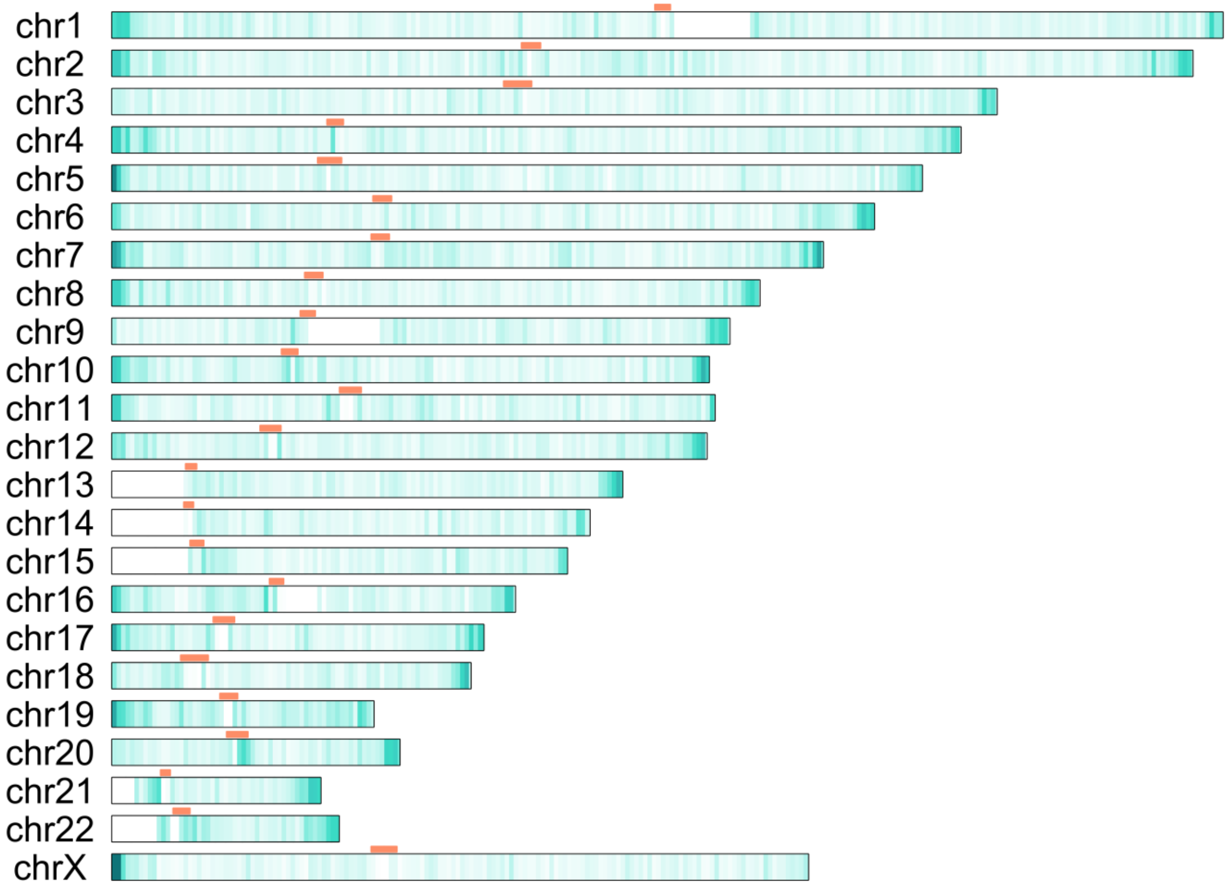

### SV overlap with functional elements

Logistic regression models were fit to determine whether SVs in high LD with GWAS SNPs have a different probability of overlapping regulatory features. The outcome regulatory feature varied, with separate models fit for open chromatin, CpG island, enhancer, promoter, TF binding, and CTCF binding overlap. The primary exposure of interest was an indicator of whether an SV had a higher LD measure with a GWAS SNP (e.g., D'≥0.8 with any nearby SNP). Since most SVs were multiallelic and LD calculations were based on binarization, there is limited interpretability of a precise LD measure such as D' or $r^2$ between an SV and a SNP. Because of this, different threshold cutoffs of D' or $r^2$ were investigated (from 0.2 to 0.8), taking note of associations that were continually significant. It is hypothesized that association between SV-SNP LD and regulatory feature overlap would indicate potential for regulatory features to aid in interpreting observed GWAS hits. The models adjusted for SV NAF, SV size, SV type (e.g., insertion, deletion), and number of genome-wide significant SNPs annotated within 1 Mb of the SV region.

**Supplementary Table 3: Structural variants in GWAS SVatalog (n = 35732) overlapping with regulatory features.**

| Database | Genomic Feature | SV Count |
|---|---|---|
| Ensembl Biomart | Enhancer | 3908* |
| | Open chromatin | 2057* |
| | CTCF Binding Site | 1699* |
| | TF Binding | 424* |
| | Promoter | 1247* |
| UCSC Genome Browser | CpG Islands | 1892 |
| | Short Tandem Repeats | 28290 |
| | Segmental Duplications | 2815 |

* Structural variants can overlap with more than one feature, some overlap with multiple

**Supplementary Table 4: Structural variants overlapping with the boundaries of genomic regions.**

| Genomic Region | SV Count |
|---|---|
| Gene boundaries | 487 |
| Start codon | 168* |
| Stop codon | 149* |
| Whole gene(s) | 278* |
| Splice sites | 1062** |
| TD Blocks | 72* |

* Addition of these categories is above the total of unique SVs overlapping gene boundaries as structural variants can encompass more than one gene
** SVs overlapping exon/intron boundaries

**Supplementary Table 5: Frequency of unique SVs encompassing entire genes.**

| Genes | 1 | 2 | 3 | 4 | 5 | 6 | 7 | 8 | 10 | 11 | 13 | 14 | 19 | 21 |
|---|---|---|---|---|---|---|---|---|---|---|---|---|---|---|
| SVs | 275 | 122 | 36 | 19 | 10 | 4 | 9 | 3 | 3 | 1 | 1 | 2 | 1 | 1 |

**Supplementary Table 7: Frequency of SVs associated with GWAS hits, using the max LD score for each SV.**

| LD stat | Range | Count | Proportion |
|---|---|---|---|
| **$r^2$** | **< 0.6** | 31947 | 89.4% |
| | **0.6 - 0.8** | 1991 | 5.6% |
| | **0.8 - 1** | 1632 | 4.6% |
| | **1** | 162 | 0.5% |
| **D'** | **< 0.6** | 7181 | 20.1% |
| | **0.6 - 0.8** | 4974 | 13.9% |
| | **0.8 - 1** | 3542 | 9.9% |
| | **1** | 20035 | 56.1% |
| **Total** | | 35732 | |
